# Bacteriological Monitoring and Evaluation of Cleaning-disinfection of Computer-related Equipment in an Obstetric and Gynecology Hospital

**DOI:** 10.1101/784017

**Authors:** Meiling Li, Tingyan Lu, Hongwei Zhang, Shufang Chen, Xueying Mao, Li Shen, Yan Lu, Shufang Leng

## Abstract

It is already known that computer keyboards and mouses in hospitals are contaminated with different kinds of bacteria. However, the mouse pad has been neglected with regard to both research and regular cleaning and disinfection in hospitals. In our study, we monitored and evaluated the bacteriology degrees of 74 computers’ keyboards, mouses and mouse pads from six departments. The results showed that before cleaning-disinfection, the contamination rate of the mouse pad ranked second following the keyboards. *Enterococcus Faecium* was cultured from the mouse pads. The computer-related equipment in the wards and outpatient rooms were much more contaminated than that in the operating rooms. *Acinetobacter spp.* was only isolated from the doctor’s computers. After cleaning-disinfection, 4 strains of MRSA were isolated from the keyboards and the mouses, one and 3 were cultured at day 3 and day 5 after cleaning-disinfection, respectively. One strain of *Pseudomonas Aeruginosa* was isolated from the mouses at day 3 after cleaning-disinfection. These demonstrated that the bacterial contamination of the mouse pads must be as emphasized as that of the keyboards and mouses. Furthermore, It is better to clean and disinfect the computer-related equipment(keyboards, mouses, mouse pads) at least once a day.

## Introduction

On a global scale, hospital-acquired infections (HAIs) have become one of the most important causes of morbidity and mortality in medical institutions[1–5] and also threaten the safety of health-care providers[6]. According to a survey from the World Health Organization, there are approximately 1.7 million and 4.5 million HAI patients in USA and Europe, respectively, accounting for 37,000 and 100,000 deaths each year. Many pathogens, such as MRSA, VRE, *Acinetobacter, Klebsiella, Listeria, Escherichia coli, Mycobacterium tuberculosis, Pseudomonas aeruginosa* and the Noel virus, can survive on a dry object surface for several months or even a year [7–9]. Therefore, cleaning and disinfecting the high-touch object surfaces is an important measure for controlling HAIs [10].

There have been many studies emphasizing the importance of cleaning and disinfecting the computer keyboard and mouse in healthcare settings, representing an important type of high-touch object surface. One study demonstrated that 95% of keyboards in a teaching hospital had growth of one or more microorganisms, and 5% were positive for pathogens known to be associated with HAI transmission, such as *Staphylococcus aureus* and *Enterococci* [11]. Some studies showed that the keyboard or mouse was one of the most likely bacterial vehicles in the ICU and that the degree of contamination cannot be neglected [12–14]. A survey of two acute district general hospitals indicated that MRSA had been identified on computer terminals (24%), and five of the MRSA-positive terminals were from hospital A, which had a significantly higher rate of MRSA transmission than hospital B [15]. However, the mouse pads have been neglected in both research and the regular cleaning-disinfection in hospitals.

In our research, we aimed to address four issues: 1) The bacteriological characteristics of computer-related equipment, especially the mouse pad; 2) The bacteriological characteristics of computer-related equipment in different clinical departments; 3) The bacteriological characteristics of doctor’s and nurse’s computer-related equipment in the wards; 4) How often we should clean and disinfect computer-related equipment.

## Materials and methods

### Study Object Selection

chosen between October 2014 and December 2015 (Supplementary Table 1). In the wards, 1 nurse’s station computer, 1 doctor’s office computer and 1 doctor’s mobile computer from each obstetric and gynecology ward were selected randomly for testing. Five samples were collected from every surface, including before cleaning-disinfection, immediately after cleaning-disinfection and day 1, day 3 and day 5 after cleaning-disinfection.

### Sample Method

Samples were collected from the keyboard (including the Number keys, Character keys, Enter key, Shift key, and Space bar), the mouse (except the underside) and the mouse pad (area≥100cm^2^). Sterile swabs dipped with sterile saline solution or neutralizing agent were smeared and rolled evenly back and forth five times on the surfaces. The hand-contacted part of the swabs was cut off, and the rest was put into a sampling tube containing 10 ml of sterile saline solution or neutralizing agent. All samples (947 samples) were sent to the clinical laboratory immediately.

### Bacteriology Identification

In a biological safety cabinet, the sampling tubes were shaken for 30s on an oscillator, 100μl of each sample was transferred to blood-agar culture medium plates, and the plates were cultured for 48 hours at 35°C in an incubator. Colonies were counted and identified by Gram stain, catalase test, oxidase test, plasma coagulase test, biochemical tube test or using a VITEK-2 instrument for bacterial identification. Drug-sensitive testing was only used for detecting MRSA.

### Statistical analysis

SPSS 17.0 software was used for statistical analysis and used the following parameters: α= 0.05, which may be calibrated according to the specific statistical data: *α*’ = 2*α* ÷ [*R* ×(*R* – 1)], in which R was the number of sample rates that had to be compared in pairs. The contamination rate (%) was the proportion of samples with bacterial colonies >10 cfu/cm^2^.

## Results

### Bacteriological Analysis before Cleaning-disinfection

### The bacterial contamination of keyboards, mouses and mouse pads

As shown in Table 1, there were significant differences in the contamination rate between the keyboard group and the mouse group, as well as between the mouse group and mouse pad group, from high to low was keyboards, mouse pads and mice, respectively. The potentially pathogenic bacteria cultured from the computer-related equipment was as shown in Supplementary Table 2, One isolate of *Enterococcus Faecium* was cultured from the mouse pad. *Klebsiella. Pneumoniae, Pseudomonas*, and *Enterobacter cloacae* were isolated from the keyboard. *Acinetobacter lwoffii* were mainly cultured from the mouse pad and keyboard.

**Table 1.** The contamination rates of keyboard, mouse and mouse pad.

|  | N | Contamination Rate (%) | Median | IQR | P-value<sup>1</sup> |
| --- | --- | --- | --- | --- | --- |
| Keyboard | 74 | 39.1 | 9.0 | 1.8-18.3 | **0.000**<sup>2</sup> |
| Mouse | 74 | 12.2 | 2.5 | 1.0-6.0 | **0.004**<sup>3</sup> |
| Mouse pad | 47 | 34.0 | 5.0 | 2.0-16.0 | 0.568<sup>4</sup> |

<sup>1</sup> P-value was calculated for the contamination rate, $\alpha' = 0.017$
<sup>2</sup> P-value was calculated between the Keyboard group and Mouse group
<sup>3</sup> P-value was calculated between the Mouse group and Mouse pad group
<sup>4</sup> P-value was calculated between the Keyboard group and Mouse pad group

### The bacterial contamination of the computer-related equipment in different departments

As shown in Table 2, the computer-related equipment in the wards and outpatient rooms were much more contaminated than that in the other departments. In total, 8 isolates of *Staphylococcus aureus* were cultured, 5, 1, 1, and 1 from the wards, outpatient rooms, neonatal dept., and delivery room, respectively. *Enterobacter cloacae*, and *Pseudomonas* were cultured from the wards. *Enterococcus Faecium* was from the neonatal dept. *Klebsiella. Pneumoniae* was isolated from the operating rooms (Supplementary Table 3).

**Table 2.** The contamination rate of computer-related equipment in different departments before cleaning-disinfection

|  | N | <sup>1</sup> Contamination Rate (%) | Median | IQR | P-value |
| --- | --- | --- | --- | --- | --- |
| Wards | 72 | 38.9 | 8.5 | 3.0-17.5 | **0.000** |
| Outpatient Room | 42 | 33.3 | 5.5 | 2.0-14.5 | **0.001** |
| Delivery Room | 14 | 28.6 | 4.0 | 1.0-15.3 | 0.036 |
| Medical Dept. | 10 | 30.0 | 3.0 | 2.0-15.0 | 0.051 |
| Neonatal Dept. | 20 | 15.0 | 0.0 | 0.0-5.8 | 0.325 |
| Operating Room<sup>2</sup> | 39 | 5.1 | 1.0 | 0.0-4.0 |  |

<sup>1</sup> Contamination rate is the subject of *P*-value calculation, $\alpha' = 0.005$
<sup>2</sup> Operating Room as the control group

### The bacterial contamination of the doctor’s and nurse’s computer-related equipment in the obstetric and gynecology wards

There was no significant difference in the contamination rate between the doctor’s office/mobile computer-related equipment and the nurse’s computer-related equipment in the obstetric and gynecology wards (Table 3). The species of potentially pathogenic bacteria from the doctor’s computers was more than that from the nurse’s computers, *Acinetobacter lwoffii* and *Acinetobacter ursingii* were isolated from the doctor’s computers. One strain of *Enterobacter cloacae* was from the nurse’s computers in the gynecology wards, 2 isolated of *Pseudomonas* were cultured from the obstetric wards (Supplementary Table 4-5).

**Table 3.**
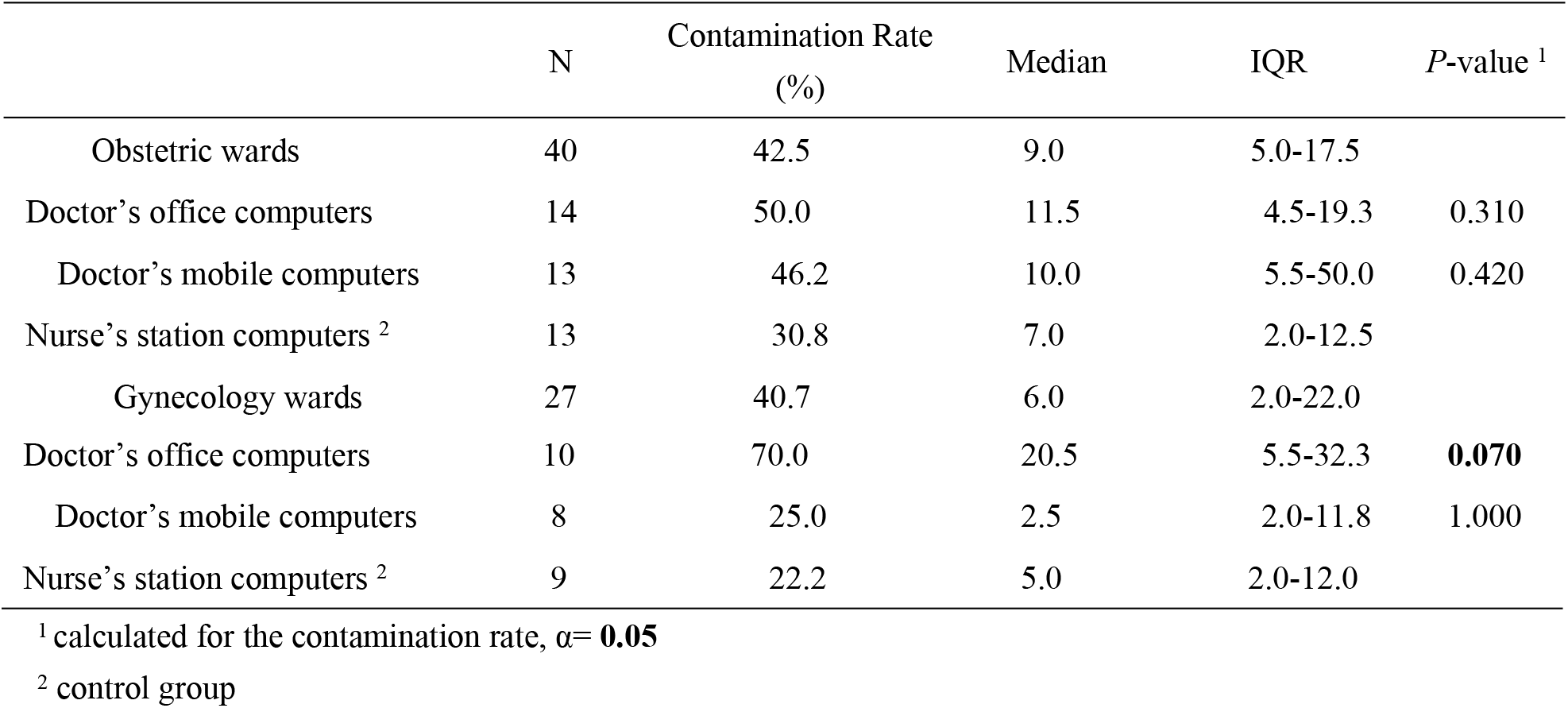
The contamination rate of computer-related equipment in obstetric and gynecology wards before cleaning-disinfection

### Bacteriological Analysis after Cleaning-disinfection

As shown in Table 4, at day 1 and day 3 after cleaning-disinfection, the contamination rates of the computer-related equipment gradually increased, and the contamination rate of mouse pads ranked the second following the keyboards. 4 strains of MRSA were isolated from the keyboards and the mouses, one and 3 were cultured at day 3 and day 5 after cleaning-disinfection, respectively. Furthermore, the strain of *Staphylococcus aureus* gradually increased (7, 8 and 10 strains).11, 7, and 7 were isolated from the keyboards, and mouses and mouse pads, respectively. One strain of *Pseudomonas Aeruginosa* was isolated from the mouses at day 3 after cleaning-disinfection (Supplementary Table 6).

**Table 4.**
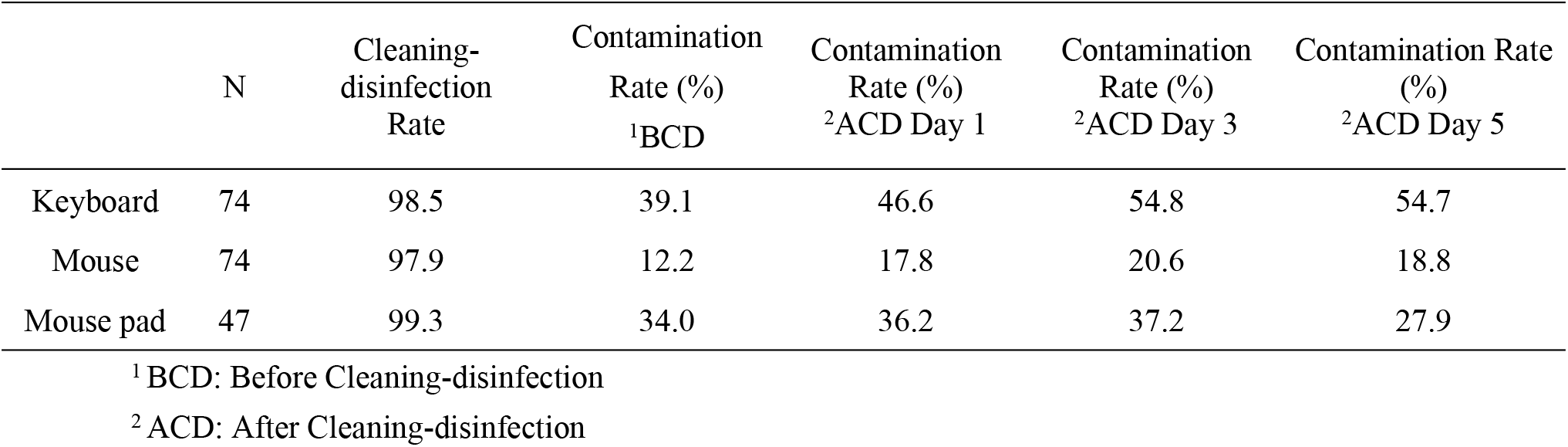
The contamination rate of computer-related equipment after cleaning-disinfection.

## Discussion

Cleaning and disinfecting object in the hospital is significantly important for controlling hospital associated infections [16–19]. In this study, we found that the contamination rate of mouse pads ranked second following the keyboards (34.0% vs 39.1%). The mouse pad is one of the high-touch objects so that it can be a “**container**” for pathogens. In another study was the contamination rate of the mouse pad researched, and the results were as same as in our study [20]. The mouse pads have been relatively disregarded in the medical settings. Furthermore, the contamination rates of computer-related equipment in the wards and outpatient rooms were significantly higher than that in the operating rooms. In the gynecology wards, the contamination rate of the doctor’s computer-related equipment was higher than that of nurse’s computer-related equipment.

The most common bacteria cultured from the computer-related equipment was *Coagulase-negative staphylococcus*, This finding was similar to the results of William’s study[21]. In total, 60 isolates of *Acinetobacter* were detected including 41 isolates of *Acinetobacter lwoffii*, 16 isolates of *Acinetobacter ursingii*, and 3 isolates of *Acinetobacter baumannii.* As an opportunistic pathogen, *A.baumannii* is one of the most clinically significant multidrug-resistant bacteria, which can cause of the nosocomial infections, especially in intensive care units [22–24]. It can persist and form biofilms on various abiotic materials in the hospital environment [24]. Contamination of ambient air with Acinetobacter baumannii was also a transmission way in Luis A study [25]. Despite *Acinetobacter spp. (A. lwoffii, A. ursingii)* other than *A. baumannii* were often considered relatively avirulent bacteria, they were able to be the opportunists in the presence of indwelling medical devices and caused invasive diseases [26]. A former research found that indwelling catheter-related with *A. lwoffii* bacteremia in immunocompromised hosts appeared to be associated with a low risk of mortality [27]. A bacteremia caused by *A. ursingii* in a patient with a pulmonary adenocarcinoma confirmed that it was an opportunistic human pathogen for the first time [28]. 33 strains of *Staphylococcus aureus* were detected, including 4 strains of MRSA. MRSA was previously detected from healthcare personnel computers [15, 29]. The other importantly isolated bacteria included *Enterococcus Faecalis, Enterococcus Faecium, Klebsiella. Pneumoniae, Enterobacter cloacae*, and *Pseudomonas Aeruginosa*.

The above isolated potentially pathogenic bacteria were also cultured from the samples of the HAI patients in our hospital, their detection rates were as shown in Table 5. the most common pathogens from HAI patients were *Enterococcus Faecalis.* this may be associated with the characteristic of maternity hospitals, with the large number of samples taken from the genital tract. *Staphylococcus aureus* and *Coagulase-negative staphylococcus* were also the main pathogens from HAI patients. The majority of *Staphylococcus aureus* were cultured from surgical incisions, and 12 cases were MRSA positive. In the process of “**patient-object-patient**” pathogens transmission, hand carriage plays an important role.

**Table 5.** The strain and percentage of the associated pathogenic bacteria from HAI patients in 2014-2016

|  | Strain (%) |
| --- | --- |
| Enterococcus Faecalis | 113 (14.9) |
| Staphylococcus aureus | 38 (5.0) |
| MRSA | 12 (1.6) |
| Coagulase-negative staphylococcus | 31 (4.1) |
| Klebsiella. pneumoniae | 26 (3.4) |
| Enterococcus Faecium | 25 (3.3) |
| Enterobacter cloacae | 9 (1.2) |
| Pseudomonas Aeruginosa | 9 (1.2) |
| Acinetobacter baumannii | 3 (0.4) |
| Acinetobacter lwoffii | 1 (0.1) |
| Micrococcus | 1 (0.1) |

A limitation of the study was the absence of bacteria homology detection among different computer equipment or between the computer equipment and HAI patients who were infected with the same bacteria. It will be further explored in the future study.

## Conclusions

In summary, we found that the cleaning and disinfection of mouse pads must be brought to attention in the hospitals. Furthermore, it was better to clean and disinfect the computer-related equipment at least 1 time/day. At the same time, health-care workers should stick to good hand hygiene.

## Acknowledgements

Not applicable

## Authors’ contributions

Meiling Li designed the study, finished all experimental tests, collected and analyzed the data and wrote the manuscript. Shufang Chen made a contribution to design the study, analyze data and wrote the manuscript. Tingyan Lu guided the bacteria test and data analysis. Hongwei Zhang contributed to guide the data analysis and modify the manuscript. Xueying Mao done a great job to revise and polish the manuscript in English. Li Shen took part in doing the experimental test. Yan Lu took part in the design of this study. Shufang Leng took part in collect the data. All authors read and approved the final manuscript.

